# Habits without Values

**DOI:** 10.1101/067603

**Authors:** Kevin J. Miller, Amitai Shenhav, Elliot A. Ludvig

**Affiliations:** Princeton Neuroscience Institute, Princeton University, Princeton, NJ, USA; Department of Cognitive, Linguistic, and Psychological Sciences, Brown Institute for Brain Science, Brown University, Providence, RI, USA; Department of Psychology, University of Warwick, Coventry, UK

## Abstract

Habits form a crucial component of behavior. In recent years, key computational models have conceptualized habits as arising from model-free reinforcement learning (RL) mechanisms, which typically select between available actions based on the future value expected to result from each. Traditionally, however, habits have been understood as behaviors that can be triggered directly by a stimulus, without requiring the animal to evaluate expected outcomes. Here, we develop a computational model instantiating this traditional view, in which habits develop through the direct strengthening of recently taken actions rather than through the encoding of outcomes. We demonstrate that this model accounts for key behavioral manifestations of habits, including insensitivity to outcome devaluation and contingency degradation, as well as the effects of reinforcement schedule on the rate of habit formation. The model also explains the prevalent observation of perseveration in repeated-choice tasks as an additional behavioral manifestation of the habit system. We suggest that mapping habitual behaviors onto value-free mechanisms provides a parsimonious account of existing behavioral and neural data. This mapping may provide a new foundation for building robust and comprehensive models of the interaction of habits with other, more goal-directed types of behaviors and help to better guide research into the neural mechanisms underlying control of instrumental behavior more generally.

## Introduction

A critical distinction exists between behaviors that are directed toward goals and those that are habitual. A large and growing body of work indicates that these behaviors depend on different sets of computations and distinct underlying neural circuits, suggesting that separable goal-directed and habitual systems implement fundamentally different strategies for the control of behavior (Balleine & O’Doherty, 2010; Dickinson, 1985; Dolan & Dayan, 2013; Wood & Rünger, 2016; Yin & Knowlton, 2006). Goal-directed behaviors are understood to be driven by consideration of the outcomes that they are likely to bring about (i.e. “action-outcome” representations). Habits, on the other hand, are understood to be driven by direct links between cues in the environment and the actions that have often followed those cues (i.e. “stimulus-response” associations). The same action may be taken under either goal-directed or habitual control in different circumstances: For example, you may take a left turn at an intersection because you have determined that turning left will get you home fastest given the specific layout of the roads and other relevant circumstances (goal-directed) or because that is what you have always done in the past at that intersection (habitual).

Several factors determine one’s likelihood of engaging in one type of behavior or another. First, habits only arise in familiar contexts, typically developing out of behaviors initially undertaken under goal-directed control (i.e., based on expected reward; Wood & Neal, 2007). Second, habits tend to form most strongly in circumstances where actions are repeated very consistently (Dickinson, 1985; Wood & Rünger, 2016). From simple motor actions to choices of meals, travel routes and exercise routines, a large body of research has demonstrated that behaviors become more automatic (i.e.., are faster, more accurate, and less susceptible to interference) the more often those behaviors are performed in the presence of a particular set of cues (reviewed in Wood & Neal, 2007; Wood & Runger, 2016). Third, the nature of one’s environment determines whether and how quickly a habit forms. Habits form slowly in environments where different behaviors lead to very different outcomes (Adams, 1982) and quickly when the environment is relatively unpredictable (Derusso et al., 2010) or when behavior is repeated with a high rate of consistency (Lally, van Jaarsveld, Potts, & Wardle, 2010). Once a behavior has become a habit, that behavior is rendered inflexible with respect to changes in the environment, including those which make the behavior undesirable (Adams & Dickinson, 1981; Hammond, 1980).

Together, these findings support a traditional view of the role of habits in instrumental control, in which they result from direct (e.g., Hebbian) strengthening of stimulus-response associations (Figure 1, left). In contrast to this view, modern computational accounts typically model habits as mediated by reinforcement-learning mechanisms, which are outcome-sensitive (Figure 1, right). Here, we argue that such computational accounts stand in tension with key data on the psychology and neuroscience of habits. We introduce a novel computational account instantiating the traditional view of habits and argue that this account provides a more parsimonious explanation for the behavioral and neural data.

**Figure 1.**
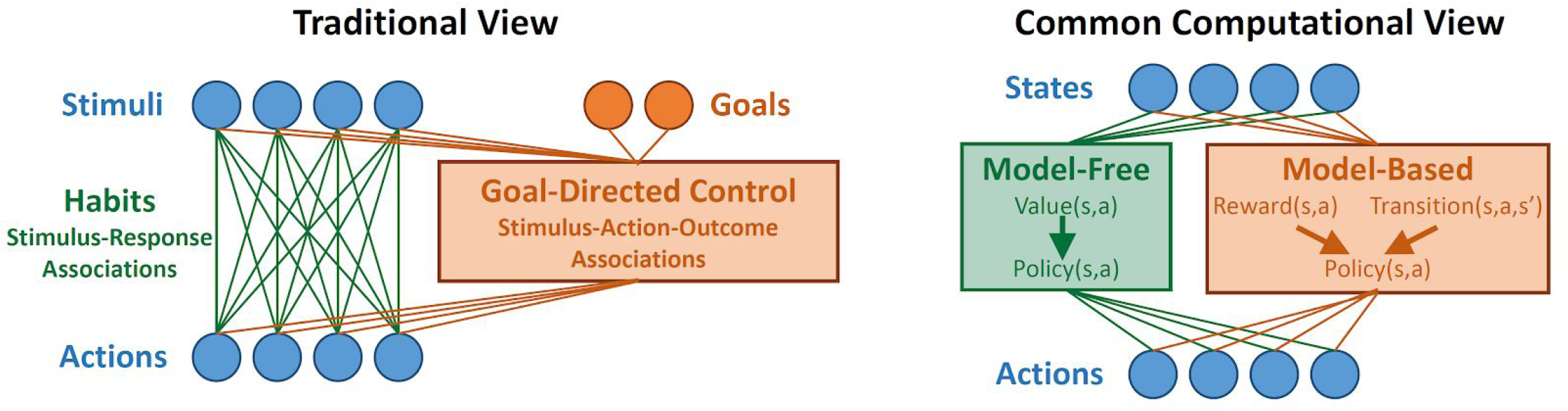
Left: Traditional view of the relationship between habits and goal-directed control. Habits are viewed as stimulus-response associations that become stronger with use, while goal-directed control takes into account knowledge of action-outcome relationships as well as current goals in order to guide choice. Right: Common computational view. Habits are implemented by a model-free RL agent which learns a value function over states and actions, while goal-directed control is implemented by a model-based RL agent which learns about the structure of the environment.

Popular computational models of habits commonly appeal to Thorndike’s “Law of Effect,” which holds that an action that has been followed by rewarding outcomes is likely to be repeated in the future (Thorndike, 1911). Modern reinforcement learning (RL) has elaborated this law into a set of computational algorithms, according to which actions are selected based on cached values learned from previous experience (Daw, Niv, & Dayan, 2005; Dolan & Dayan, 2013; Sutton & Barto, 1998). This class of computations focuses only on potential rewards, ignoring all reward-unrelated elements of one’s environment (discussed below); it is therefore referred to as “model-free” RL. This formulation for habits has become so prevalent that the terms “habit” and “model-free” are now used interchangeably in much of the computational literature (Dolan & Dayan, 2013; Doll, Simon, & Daw, 2012). Equating these terms, however, carries a critical assumption: that habits are driven by a reward-maximization process (i.e., a process that depends directly on potential outcomes).

Model-free algorithms typically operate by learning the expected future reward associated with each possible action or state, relying crucially on these value representations. This idea, that habits are *value-based*, strains against traditional interpretations of habits as stimulus-response (S-R) associations that are blind to potential outcomes (Dickinson, 1985; Hull, 1943; James, 1890). The latter, *value-free* definition for habits drove the development of critical assays that have been used to discriminate between actions that are habitual versus goal-directed (i.e., outcome-sensitive), testing whether an animal continues to pursue a previous course of action when that action is no longer the most beneficial (Adams & Dickinson, 1981; Hammond, 1980). Such a value-free formulation of habits aligns well with Thorndike’s second law, the “Law of Exercise.” This law holds that an action that has been taken often in the past is likely to be repeated in the future, independent of its past consequences; in other words, it describes habits as a form of perseveration. This category of value-free habits has been maintained in modern theorizing on habits, where it has been referred to as “direct cuing” of behavior, as distinct from value-based forms of habits (“motivated cueing”; Wood & Neal, 2007; Wood & Rünger, 2016).

Here, we develop a computational implementation of the Law of Exercise and show that it offers an alternative to model-free RL as a mechanism for habits, one that retains ideas about the nature of habits that have developed within other areas of psychology and neuroscience (Graybiel, 2008; James, 1890; Wood & Rünger, 2016). This value-free habit mechanism accounts for key findings in the animal learning literature that dissociate habitual and goal-directed actions, namely the tendency for an animal to continue performing a previously learned action when that action is no longer predictive of the reinforcing outcome (contingency degradation; Hammond, 1980) or when the predicted outcome ceases to be desired by the subject (outcome devaluation; Adams & Dickinson, 1981). In addition, this model provides what is, to our knowledge, the first computational account of the difference in rate of habit formation under variable-interval and variable-ratio schedules of reinforcement (Adams, 1982; Dickinson, 1985; Gremel & Costa, 2013). Furthermore, a value-free habit mechanism explains a variety of other behavioral phenomena in which responses are facilitated by simple repetition (i.e. perseveration Aarts, Verplanken, & van Knippenberg, 1998; Akaishi, Umeda, Nagase, & Sakai, 2014; Akam et al., 2017; Balcarras, Ardid, Kaping, Everling, & Womelsdorf, 2016; Bertelson, 1965; Cho et al., 2002; Gold, Law, Connolly, & Bennur, 2008; Gore, Dorris, & Munoz, 2002; Jung & Dorner, 2018; Kim, Sul, Huh, Lee, & Jung, 2009; Lau & Glimcher, 2005; D. Lee, McGreevy, & Barraclough, 2005; Riefer, Prior, Blair, Pavey, & Love, 2017)

In addition to reformulating the computational underpinnings of habits, we will show that our model offers a critical realignment to prevailing models of instrumental control. According to this prevailing computational framework (Figure 1, right), an equivalence between habitual control and model-free RL computations is paralleled by an equivalence between goal-directed behavior and another set of RL computations, referred to as “model-based” RL (Daw et al., 2005; Dolan & Dayan, 2013). Model-based RL guides behavior through an internal model of the environment that is used to estimate values for each action. This internal model of the environment includes both the expected likelihood of transitioning between environmental states and the expected rewards for each action. A substantial theoretical and empirical literature has been built around the idea that the habitual/goal-directed distinction can be equated with the model-free/model-based distinction from RL, and this presumed equivalence has been used to glean insights into complex decision-making phenomena, such as addiction (Lucantonio, Caprioli, & Schoenbaum, 2014; Redish, Jensen, Johnson, & Kurth-Nelson, 2007), impulsivity (Kurth-Nelson, Bickel, & Redish, 2012; Rangel, 2013), compulsivity (Gillan, Kosinski, Whelan, Phelps, & Daw, 2016; Gillan, Otto, Phelps, & Daw, 2015), and moral judgement (Buckholtz, 2015; Crockett, 2013; Cushman, 2013). By replacing model-free RL with a value-free mechanism, our model forces a critical realignment of this prevailing framework, thereby prompting a deeper consideration of how the computations and circuitry for model-free and model-based reinforcement learning might share more commonalities than differences.

## Methods

### Computational Model

As proof of concept, we implemented the proposed mechanisms for habitual and goal-directed control in a computational model. This model contains three modules: a goal-directed controller, a habitual controller, and an arbiter (Figure 2). The goal-directed controller is sensitive to outcomes, selecting actions that are likely to lead to outcomes that have high value. Here, we instantiate it using a model-based reinforcement learning algorithm. The habitual controller, on the other hand, is sensitive only to the history of selected actions. It tends to repeat actions that have frequently been taken in the past (e.g., because they were selected by the goal-directed controller), regardless of their outcomes. The arbiter weights the influence of each of these controllers on behavior, tending to favor goal-directed control when action-outcome contingency is high, and to favor habitual control when habits are strong.

**Figure 2:**
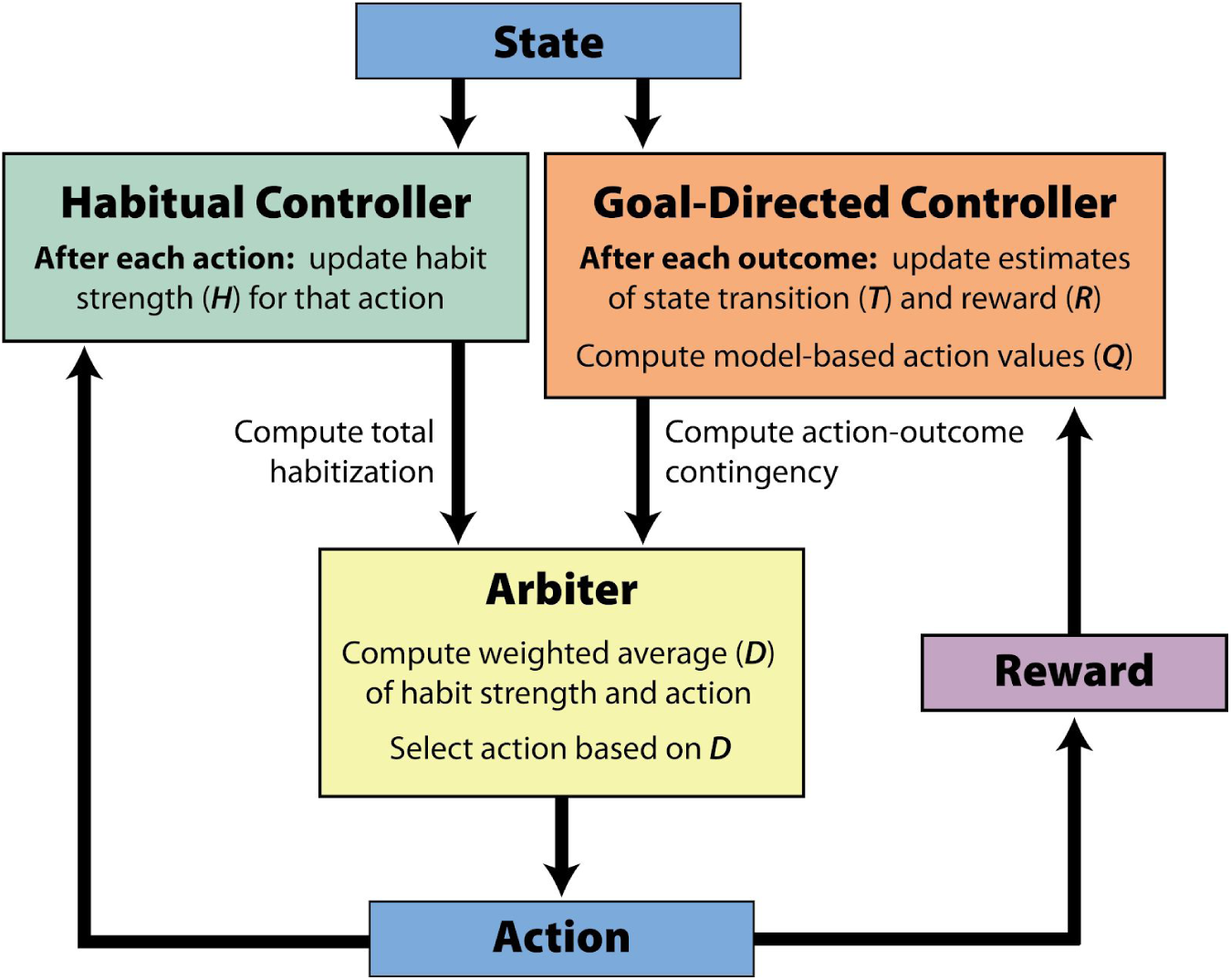
Schematic description of the model components and their interactions. See main text for details.

#### Habitual Controller

The habitual controller is sensitive only to the history of selected actions, and not to the outcomes of those actions. This action history is tracked by a matrix of habit strengths, **H**_**t**_, in which *H*_*t*_(s,a) acts as a recency-weighted average of how often action *a* was taken in state *s* prior to timepoint *t*. Initial habit strength **H**_**0**_ is set to zero and updated after each trial according to the following equation:

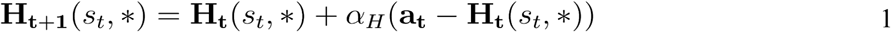

where *s*_*t*_ is the current state, **H**_**t**_(s_t_,*) is the row of **H**_**t**_ corresponding to *s*_*t*_, *α*_*H*_ is a step-size parameter that determines the rate of change, and **a**_***t***_ is a row vector over actions in which all elements are zero except for the one corresponding to *a*_*t*_, the action taken on trial *t*. Note that the particular environments simulated in this paper all include only a single state, so for this and all subsequent equations we will drop the indexing by *s*, and consider **H** to be a vector over actions:

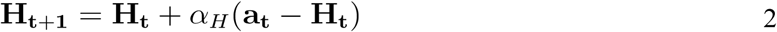

For a full version of the model suitable for environments with multiple states, see equations in Appendix A.

#### Goal-Directed Controller

The goal-directed controller is composed of a model-based RL agent, sensitive not only to the actions taken, but also to their outcomes. In contrast to traditional reinforcement-learning methods, this agent does not consider “common currency” reward, but rather learns separately about reinforcers of different types. It maintains an estimate, **R**_**t**_ of predicted immediate reinforcement, in which *R*_*t*_*(a,m)* gives the agent’s expectation at timepoint *t* of the magnitude of reinforcer type *m*, that will follow from action *a*. Initial reinforcement expectation **R**_**0**_ is set to zero, and after each trial, the agent updates these quantities according to the following equations (Sutton & Barto, 1998):

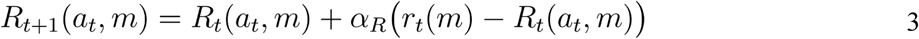

where *a*_*t*_ is the current action, *r*_*t*_*(m)* is the magnitude of the reinforcer of type *m* received following that action, and *α*_*R*_ is a step-size parameter which governs the rate of learning. The full model, suitable for environments with multiple states, includes equations for learning about the state transitions, as well as for estimating the expected future value associated with each action using planning (see Appendix A). In environments with only one state, the expected value for each action *Q(a)* is based on the expected immediate reinforcement of each type, as well as the agent’s utility for reinforcers of each type:

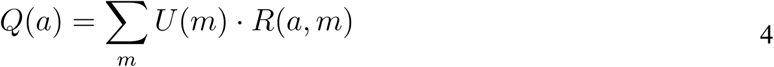

where *U(m)* is a utility function giving the value that the agent assigns to reinforcers of each type *m*. This value is typically unity for reinforcers designated “food pellets”, 0.1 for reinforcers designated “leisure”, and −1 for reinforcers designated “effort”, unless otherwise noted (see *Simulation Three: Outcome Devaluation*).

#### Arbiter

The arbiter governs the relative influence of each controller on each trial. It computes an overall drive *D(a)* in favor of each action, *a*, as a weighted sum of the habit strength *H(a)* and the goal-directed value *Q(a)*.

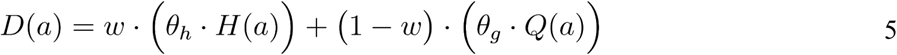

Where *θ*_*h*_, and *θ*_*g*_ are scaling parameters, and *w* is a weight computed on each trial by the arbiter to determine the relative influence of each controller (see Equation 9). The model then selects actions according to a softmax on **D**:

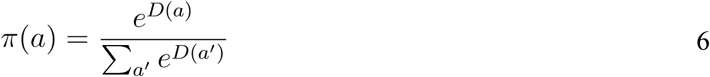

To determine the appropriate weight *w*, the arbiter computes two quantities, the *action-outcome contingency* (*g*) and the overall *habitization* (*h*), which promote goal-directed and habitual control, respectively. Action-outcome contingency is a measure of the extent to which the expected outcome received varies according to the action that is performed. Here, we quantify action-outcome contingency for a particular reinforcer *m*, conditional on a particular policy ***π*** with the following equation:

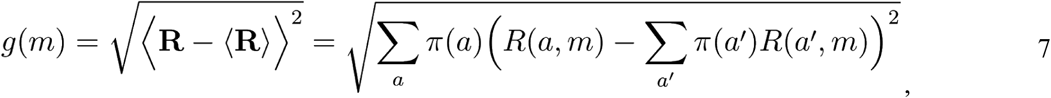

which reflects the degree of variation in expected outcome for that reinforcer, based on the available actions and the policy. The measure *g* is minimal when all actions have the same expected outcome and increases as the outcomes associated with some actions are increasingly distinct from the outcomes associated with other actions. Note that this measure considers the degree to which *average expected* outcome varies with action. It does not consider the degree to which *particular outcomes* vary conditional on particular actions (e.g., an environment in which one action led to one pellet with certainty and another led to two pellets with 0.5 probability would be rated as having zero action-outcome contingency because the average outcome is identical for both actions). In our simulations, we include two types of reinforcers: “food pellets” and “leisure”. Because leisure is not a true outcome in the environment, we compute *g(m)* with respect to food pellets only, and drop the index by reinforcer type, considering *g* to be a scalar in future equations. The arbiter also computes an analogous quantity for the habitual controller, which we term “overall habitization” *h*:

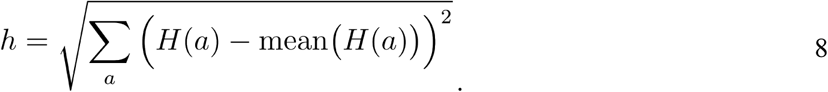

The overall habitization *h* is minimal when no action has a large habit strength, or when all action have approximately equal habit strengths. It is maximized when one or a few actions have much larger habits strengths than the others. The arbiter then computes the mixing weight *w* on the basis of these two quantities:

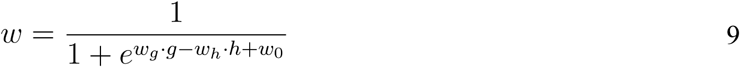

where *w*_*g*_ and *w*_*h*_ are scaling parameters controlling the relative strengths of the goal-directed and habitual systems, w_0_ is a bias parameter, which shifts control toward the goal-directed system. Note how *g* is no longer dependent on reinforcer type because all simulations in this paper contain only one type of reinforcer other than leisure. This calculation represents a push-pull relationship whereby goal-directed control is facilitated to the extent that the action-outcome contingency is high, whereas habits are facilitated to the extent that habitization is large. Figure 3 provides an intuition for how each of the values described above evolves in a setting where the more valuable of two actions reverses at some point in a session.

### Simulated Task Environments

#### Simulation 1: Reversal Learning

To illustrate the behavior of the model and the dynamics of its various internal variables, we simulated behavior in a probabilistic reversal learning task (Fig 3). In this task, the agent was presented with an environment consisting of a single state in which two actions were available. In the first phase of the task (1000 trials), performance of one action (Action A) resulted in a reinforcer 50% of the time, while performance of Action B never did. In the second phase (reversal), Action A never resulted in a reinforcer, while Action B resulted in one 50% of the time that it was taken.

**Figure 3.**
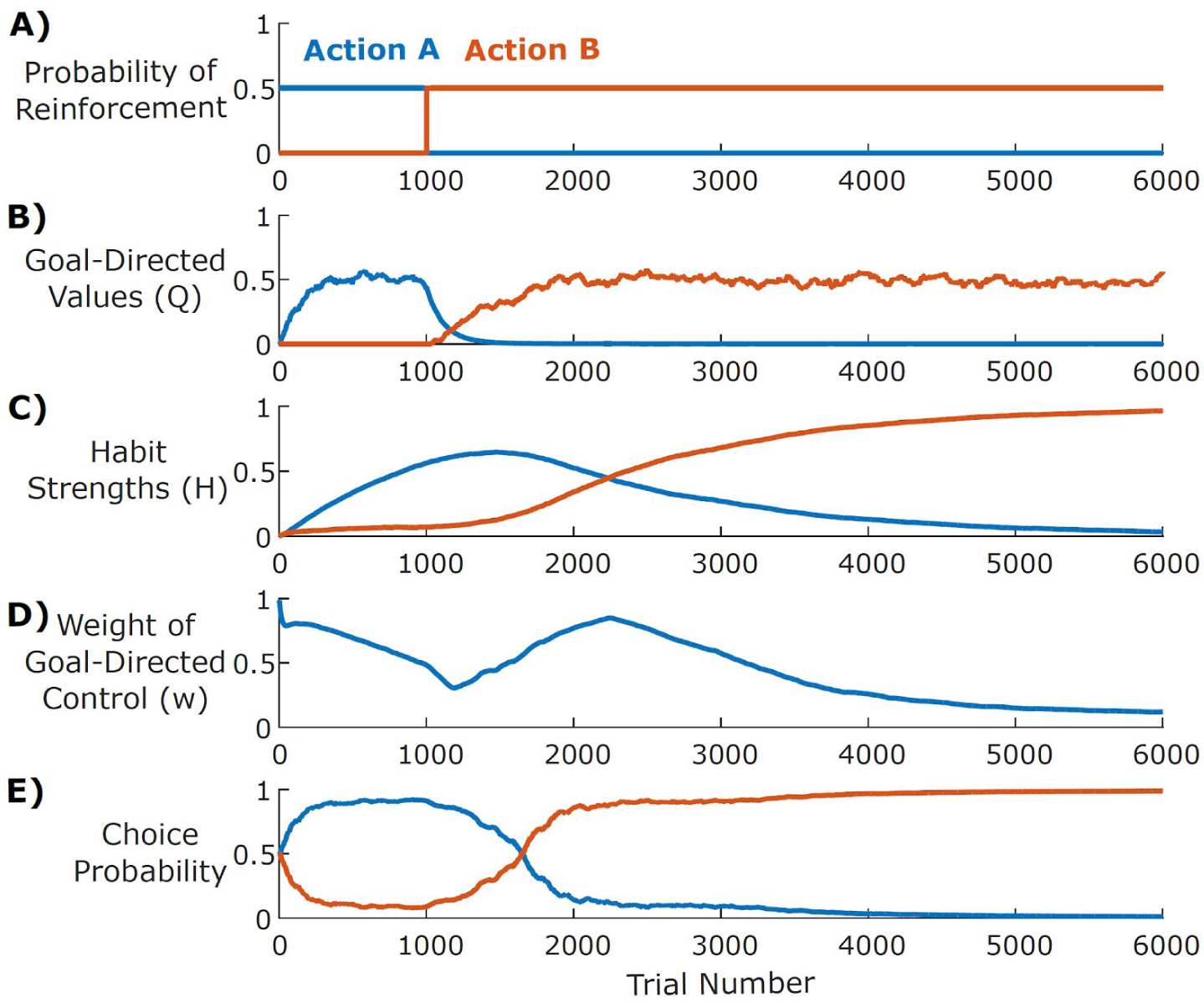
A) Simulations of a reversal-learning environment: Action A is initially reinforced with higher probability (0.5) than Action B (0), but after 1000 trials, the relative dominance of the actions reverses. B) Soon after the reversal, the goal-directed system learns that Action B is more valuable. C) The habit system increasingly favors Action A the more often it is chosen and only begins to favor Action B once that action is chosen more consistently (long after reversal). D) The weight of the goal-directed controller gradually decreases as habits strengthen, then increases post-reversal as the global and goal-directed reinforcement rates diverge. E) Actions are selected on each trial by a weighted combination of the goal-directed values (*Q*) and the habit strengths (*H*) according to the weight (*w*).

#### Simulation 2: Omission Contingency

We simulated behavior in an omission experiment using a similar environment with one state and two available actions. In the first phase, performance of one action (Press Lever) resulted in a reinforcer of one type (Pellet) 25% of the time, while performance of the other action (Withhold Press) resulted in a reinforcer of another type (Leisure) 100% of the time. In the second phase, performance of Press Lever was never reinforced, but performance of Withhold Press resulted in both a 25% chance of Pellet and a 100% chance of Leisure. We set the agent’s utilities for Pellet and Leisure to 1.0 and 0.1, respectively. To investigate the effect of training duration on behavioral flexibility in the face of omission, we varied the number of trials in the training phase from 100 to 2000, in intervals of 100. The omission phase was always 500 trials in duration. We simulated ten agents for each duration of phase one, and report the average rate of performance of Press Lever for the final trial in each phase.

#### Simulation 3: Outcome Devaluation

We simulated behavior in an outcome devaluation experiment in a similar way. The training phase was identical to that used for omission. This training phase was followed by a devaluation manipulation, in which we set the agent’s utility for the Pellet reinforcer to 0, and then an extinction phase. In the extinction phase, performance of Press Lever resulted in no outcome, while performance of Withhold Press continued to result in Leisure with probability 100%. To investigate the effect of training duration on behavioral flexibility in the face of devaluation, we again varied the number of trials in the training phase from 10 to 2000 in intervals of 100, and report the average rate of performance of Press Lever for the final trial in each phase.

#### Simulation 4: Framework for Free-Operant Tasks

Assays of goal-directed and habitual behavior are typically performed not in two-alternative forced-choice environments like those we describe above, but rather in free-operant tasks in which subjects are not constrained by a discrete-trial structure, but are free to perform one or more actions (e.g., lever presses) at any rate they wish. To simulate these environments, we adjusted our model to accommodate choices along this continuous variable (lever press rate). In this simulation, the actions (*a*) were press rates, which ranged between 0 and 150 presses per minute. This extension to a larger action space required two changes to the model. The first was the use of function approximation to compute the reinforcer function **R** and the habit strength **H**. Instead of learning these functions directly over each value of *a*, as in equation 2 or 3, the model approximated them using a set of basis functions. For the reinforcement function, we used a Taylor (polynomial) basis set with four bases:

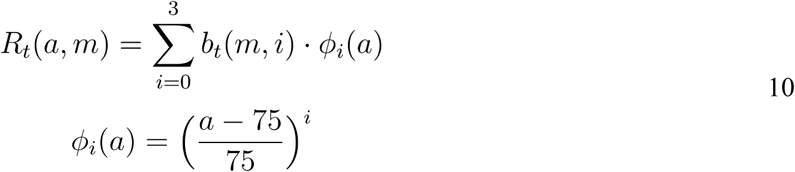

where *b*_*t*_*(m, i)* is the learned weight for each reinforcer *m* and basis element *i* (see Equation 12). For the habit strength function *H(a)*, we used a set of radial basis functions, with 30 Gaussian bumps with means at intervals of 5 and standard deviation 5 presses per minute:

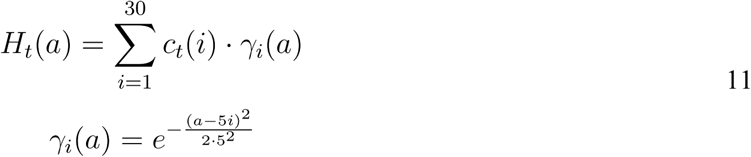

where *c*_*t*_(i) is the learned weight for corresponding basis element *i*. The goal-directed weights *b* and the habitual weights *c* are then updated using a stochastic gradient descent procedure (Sutton & Barto, 1998):

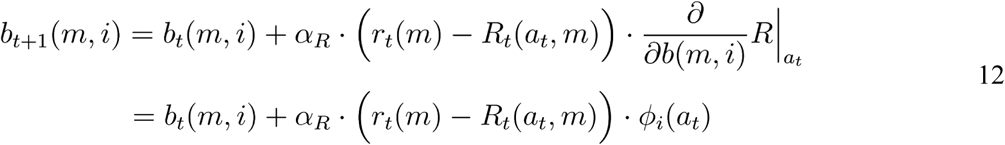

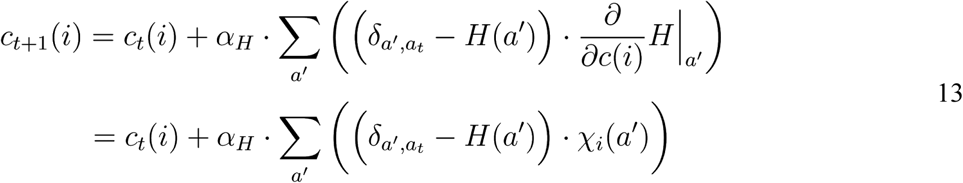

where 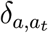 is a Kronecker delta function, taking on a value of one when *a* = *a*_*t*_ and zero otherwise, and the vertical bar with subscript indicates that the partial derivative is evaluated at the point indicated.

The second change we made was to introduce an “action density” measure *m*, which can be thought of as controlling how many distinct “actions” are available that yield a particular press rate. Selecting an action density measure that is sharply peaked at zero ensures that an agent that chooses randomly will on average press at a low rate, rather than selecting at random from a uniform distribution of press rates (i.e., pressing on average at half of the maximum possible rate). We used an exponentially decaying action density function with a scale of 5.

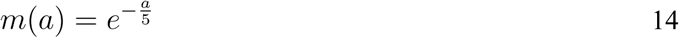

This measure influences action selection, leading to a tendency to prefer rates for which more actions are available (i.e., low press rates). In place of Equation 6, the model now selects actions according to:

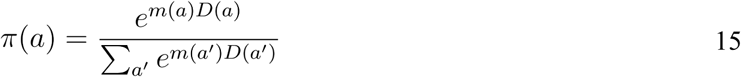

#### Habitization in Variable Interval vs. Variable Ratio Reinforcement Schedules

We used the above framework to simulate behavior under two reinforcement schedules commonly used in experiments on animal learning, termed “variable ratio” (VR) and “variable interval” (VI) schedules. In a VR schedule, the probability of receiving a reinforcer is constant after each lever press. Reinforcement rate is therefore directly proportional to response rate and potentially unbounded. In a VI schedule, reinforcers are “baited” at variable intervals, and the first press following baiting will lead to a reinforcer. The probability that a press will be reinforced therefore increases as a function of the time since the last press, and reinforcement rate is a sublinear function of response rate, saturating at the average baiting rate. In both environments, lever pressing is thought to involve some effort cost, which increases superlinearly with the rate of responding. To model acquisition in a VR environment, we used VR10, in which each press had a 10% chance of being followed by a pellet. To model acquisition in a VI environment, we used VI6, in which pellets were baited every six seconds, or on average ten times per minute. In both cases, we included an effort cost that was quadratic in press rate. Specifically, each action resulted in reinforcers of two types: Pellet Rate, with positive utility, and Effort Rate, with negative utility. Effort was modulated by press rate, to reflect the physical and cognitive costs associated with lever pressing:

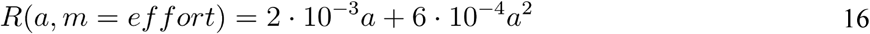

where *a* is the press rate selected, with units of presses per minute.

#### Omission and Devaluation in VR vs. VI Schedules

To investigate the effects of training duration on behavioral flexibility in these free-operant environments, we exposed agents given limited training (5,000 trials) or extended training (30,000 trials) with either a VR or a VI schedule to both Omission and Devaluation manipulations. In the Omission manipulation, we changed the reinforcement schedule such that the magnitude of the Pellet Rate reinforcer was inversely related to the Press Rate action. The magnitude of the Leisure reinforcer (reflecting effort cost) was not changed. In the Devaluation manipulation, we left the reinforcement schedule unchanged, but changed the agent’s utility for the Pellet Rate outcome to 0.

#### Lesions of Goal-Directed vs. Habitual Controllers

To simulate lesions of goal-directed and habitual controllers on behavior in free-operant tasks, we repeated the above experiments with the parameters of the model altered. Specifically, to model lesions of the goal-directed controller, we decreased the parameters *θ*_*g*_ and *W*_*g*_, whereas to model lesions to the habitual controller, we decreased the parameters *θ*_*h*_ and *W*_*h*_ (see Table 1 for details).

**Table 1.**
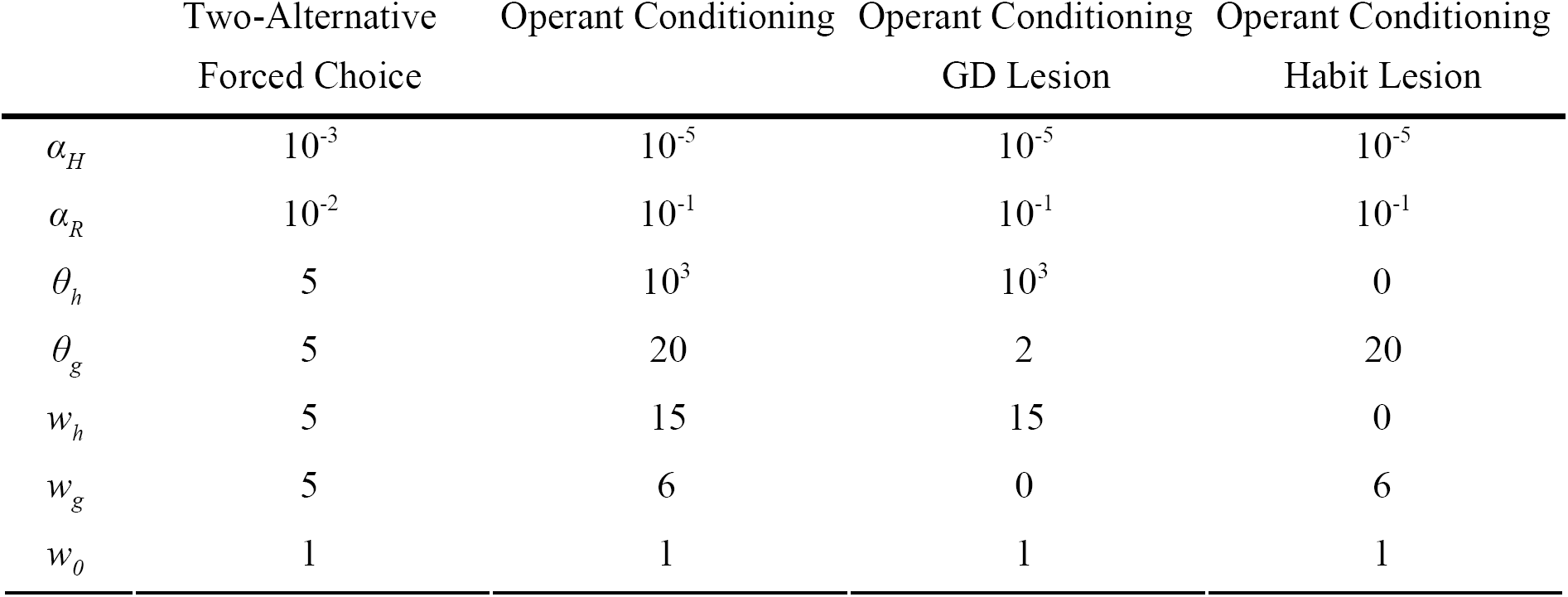
Parameter values used in simulations

#### Two-Armed Bandit Task

To illustrate the role of habits in producing perseveration in free-choice tasks, we simulated data from our agent performing a two-armed bandit task. The model was tested in an environment consisting of one state in which two actions were available. Performing either of these actions led to a reinforcer with some probability. Reinforcer probabilities were initialized uniformly between 0 and 1, and changed slowly across trials according to independent Gaussian random walks (SD = 0.15; bounded at 0 and 1), requiring the agent to continuously learn. The agent performed 10,000 trials in this environment, using the parameters in Table 1. Task parameters were selected to facilitate comparison to a rodent behavior dataset using a similar task (Miller, Botvinick, & Brody, in prep; Miller, Erlich, Kopec, Botvinick, & Brody, 2013). To simulate a dataset with similar characteristics to the rat dataset, we generated data from 50 copies of our model, with parameters sampled from the range described in Table 2. See Appendix B for a detailed description of this agent. We analyzed these datasets using a logistic regression model that quantifies the influence of previous choices and their outcomes on future choice (Lau & Glimcher, 2005; Miller et al., in prep).

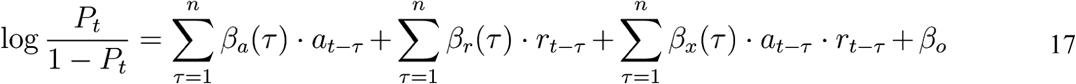

where *P*_*t*_ is the probability that the model believes the agent will select action 1 on trial *t, a*_*t*_ is the action taken on trial *t, r*_*t*_ is the reinforcer received, *n* is a parameter of the analysis governing how many past trials to consider, *β*_*a*_, *β*_*r*_, and *β*_*x*_ are vectors of length *n* containing fit parameters quantifying the influence of past actions, reinforcers, and their interaction, respectively, and *β*_*o*_ is an offset parameter. Positive fit values of *β*_*a*_ indicate a tendency of the agent to repeat actions that were taken in the past, independently of their outcomes, while positive values of *β*_*x*_ indicate a tendency to repeat actions that led to reinforcement and to switch away from actions that do not.

**Table 2.**
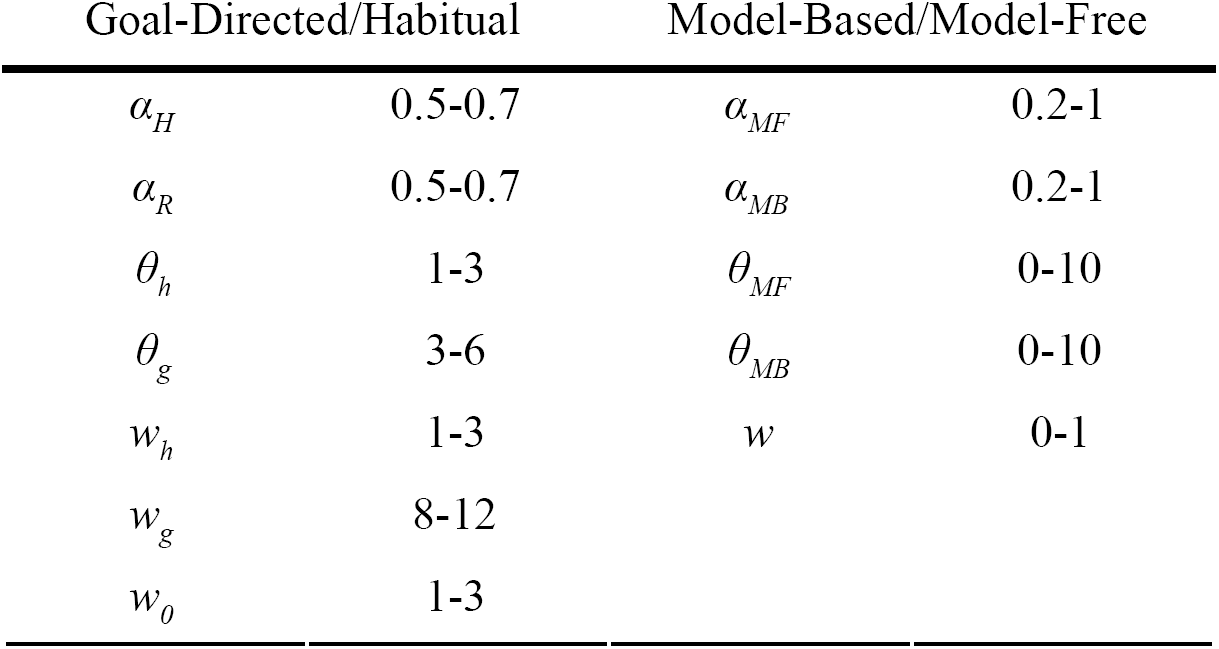
Parameter ranges used in simulations of two-armed bandit task

## Results

Our model proposes that behavior arises from the combined influence of two controllers: one driven by value-free perseveration (habitual) and one driven by model-based reinforcement learning (goal-directed). Figure 3 illustrates the learning dynamics of these two controllers in a simple reversal learning task, where an animal first learns to associate Action A with a higher probability (0.5) of reinforcement than Action B (0) and, after 1000 trials, this contingency reverses (Figure 3A). Initially, behavior is driven by the goal-directed controller, which gradually learns the relative reinforcement rates (Figure 3B) and thus increasingly selects Action A. As it does so, the habitual controller strengthens its association between the current stimuli and Action A (Figure 3C). As these habits strengthen, the habitual controller increasingly drives the choice of which action to select (Figure 3D). As a result, when the reversal occurs, the agent continues to select Action A for an extended period, past the point where the goal-directed controller has learned that Action B is more likely to be reinforced (compare Figures 3B and 3E). In addition to demonstrating the different kinds of learning that drive each of these controllers, this example demonstrates that our model captures the observation that behavioral control in a novel environment tends to evolve from goal-directed to habitual (i.e., habits form from actions that were originally selected in a value-based manner). In the following sections we demonstrate that this model can capture all of the key diagnostic features of habitual behavior previously identified in animal behavior, including sensitivity to repetition frequency, reinforcement schedule, and selective modulation by lesions to one of two dissociable neural circuits.

### Effects of Training Duration on Behavioral Flexibility

We first test whether our model can capture a central finding from research on habits: that extensive training in a static environment (overtraining) can render behavior inflexible in the face of changes to that environment. This inflexibility is classically demonstrated in two ways: by altering the contingencies between actions and outcomes, or by devaluing the outcomes themselves. The first of these manipulations involves altering the probability that an outcome (e.g., a food pellet) will be delivered following an action (e.g., a lever-press) and/or the probability that it will be delivered in the absence of the action (contingency degradation). Overtrained animals will often continue to perform an action even when it no longer causes the desired outcome (indeed, even when it *prevents* the delivery of the outcome). This perseverative behavior is diagnostic of habitual control (Dickinson, 1998). The second manipulation involves rendering the outcome no longer desirable to the animal (e.g., by pairing its consumption with physical illness) – in this setup, overtrained animals will often continue performing an action that leads to an outcome they no longer desire (Adams, 1982).

We simulated the effect of overtraining on sensitivity to an omission contingency by running the model through simulated sessions with two stages. The agent was initially trained in an environment where performance of Action A (“press lever”) was followed by a reinforcer of one type (“food pellet”) 50% of the time, and Action B (“withhold press”) was followed by a reinforcer of another type (“leisure”) 100% of the time. The agent’s utility for the food pellet reinforcer was set to 1, while the utility of the leisure reinforcer was set to 0.1, and with experience in this environment the agent learns to press the lever on a large fraction of trials (Figure 4, blue curves). After a number of trials that varied between simulations, the reinforcement probabilities for the food pellet were reversed, such that pressing the lever resulted in no reinforcement, and withholding resulted in leisure 100% of the time and a food pellet 50% of the time. When the agent was given a small number of training trials, it successfully learned to decrease probability of lever pressing following this reversal. With longer training sessions, however, the model failed to reverse its actions within the same time period (Figure 3, left). This is consistent with data from animal learning, in which overtraining abolishes behavioral sensitivity to omission contingencies (Dickinson, 1998)

**Figure 4.**
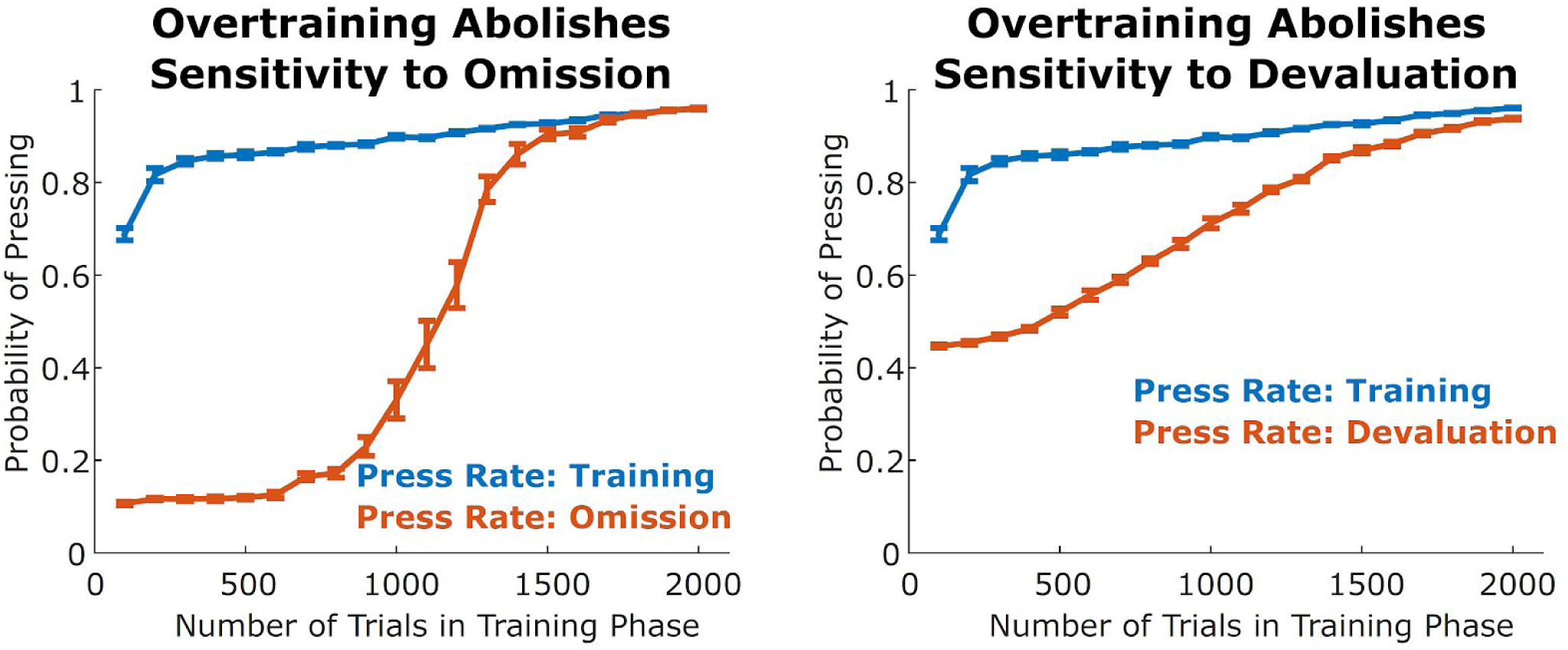
Behavior becomes inflexible after overtraining. Rate of pressing in a simulated instrumental conditioning task at the end of the training period (blue) as well as following omission or devaluation manipulations (orange), as a function of the duration of the training period. As this duration increases, the agent is increasingly unlikely to alter its behavior (blue and orange curves become similar). These simulations are consistent with the finding that overtraining results in behavior that is insensitive to omission and to devaluation. Error bars represent standard errors over ten simulations.

We simulated the effect of overtraining on outcome devaluation in a similar manner. The first stage of training was similar, with lever pressing being followed by a pellet 50% of the time and withholding being followed by leisure 100% of the time. At the end of this first stage, the agent’s utility for the food pellet reinforcer was decreased to zero, simulating a devaluation manipulation. In the final stage (testing), the agent was placed back in the choice state, and had the opportunity to again select between pressing and not pressing, with the outcome of pressing no longer delivering any reinforcement. We found that the frequency of choosing to lever-press in the testing stage strongly depended on the duration of the training stage (Figure 3, right), indicating that overtraining caused the agent to perseverate on the habitized behavior (lever-pressing).

### Effects of Reinforcement Schedule on Habit Formation

Another central finding from research on habitual control is that the reinforcement schedule has a profound effect on habit formation. Actions followed by a constant probability of reinforcement (variable-ratio [VR] schedules) take a long time to habitize (Adams, 1982). By contrast, when an action is “baited” at a constant probability per unit time, and only the first press following baiting results in reinforcement (variable-interval [VI] schedules), the action habitizes much more quickly, even when performance and overall rate of reinforcement are matched across these two reinforcement schedules (Dickinson, Nicholas, & Adams, 1983). The finding that VI schedules result in rapid habit formation has been widely replicated and represents a key element of the experimental toolkit for the study of habits (Gremel & Costa, 2013; Tanaka, Balleine, & O’Doherty, 2008; Yin & Knowlton, 2006). We sought to replicate this effect using our model.

To do this, we adapted the model to operate in a continuous action space (see Methods for details). Briefly, at each timestep, instead of making a binary decision (e.g., between pressing the lever or not pressing), the agent instead selected a scalar lever pressing *rate*. Accordingly, the agent then observed a rate for each reinforcer rather than binary reinforcement. As the action Press Rate increased, two types of reinforcement increased, one with positive utility (Pellet Rate) and the other with negative utility (Effort Rate). For VR schedules, Pellet Rate is a linear function of Press Rate (Figure 4, top right), because each press results in reinforcement with equal probability. For VI schedules, Pellet Rate is a sublinear function of Press Rate, saturating at the rate of baiting (no matter how often a rat in a VI experiment presses the lever, reinforcers are only available as they are baited; Figure 4, top left). In both schedules, we modeled Effort Rate as a superlinear function of Press Rate (see Methods). The agents used function approximation to learn estimates for these two functions (which together comprise the model for the goal-directed controller), as well as for habit learning.

Consistent with empirical findings, we found that simulated agents trained on VI schedules lever-pressed at a moderate rate and habitized early in training (Figure 4, left) whereas agents trained on VR schedules lever-pressed at a high rate and habitized much later in training (Figure 4, right). This difference was largely driven by the difference in action-outcome contingencies inherent to each schedule: a small change in press rate resulted in a much larger change in reinforcement rate for a VR schedule relative to a VI schedule (compare the slope of the green curve in the right panel relative to the left panel of Figure 4).

### Effects of Striatal Lesions on Habit Formation and Behavioral Flexibility

The degree to which an animal behaves flexibly in a given environment can be affected profoundly by manipulating specific brain structures. Lesions to regions of a putative “habit system,” such as the dorsolateral striatum (DLS), promote behavioral flexibility (i.e., alleviate perseveration) following overtraining (Graybiel, 2008; Yin & Knowlton, 2006). Conversely, lesions to regions of a putative “goal-directed system,” such as the dorsomedial striatum (DMS) impair flexibility (Yin, Ostlund, Knowlton, & Balleine, 2005). In particular, relative to control rats, rats with lesions to DMS lever-press at a lower rate and are less able to decrease their press rate when reinforcers are omitted or devalued (Yin et al., 2005); rats with lesions to DLS lever-press at a similar rate to controls and are more successful than controls at adapting their press rate to omitted or devalued reinforcement (Yin, Knowlton, & Balleine, 2004, 2006).

We lesioned the goal-directed controller (DMS) or habitual controller (DLS) in our model while simulated agents performed the free-operant task (see Table 1 for details). These agents received either limited training (5,000 trials) or extensive training (15,000) in the VR environment. They were then subjected to either an omission contingency (greater rates of lever pressing caused lower rates of reinforcement) or a devaluation manipulation (the utility of the pellet reinforcer was set to zero). Consistent with empirical findings described above, we found that lesioning the goal-directed system produced a low press rate that was unaffected by either omission or devaluation, whereas lesioning the habitual controller led to a high press rate that adapted to both manipulations (Figure 5). A “control” agent, with intact habitual and goal-directed controllers, adopted a high press rate that adapted to both manipulations when given limited training, but did not adapt to either following extensive training. Lesions to our model’s goal-directed and habitual controllers thus reproduce behavioral patterns typical of DMS and DLS lesioned rats, respectively, in classic experiments on instrumental conditioning.

**Figure 5.**
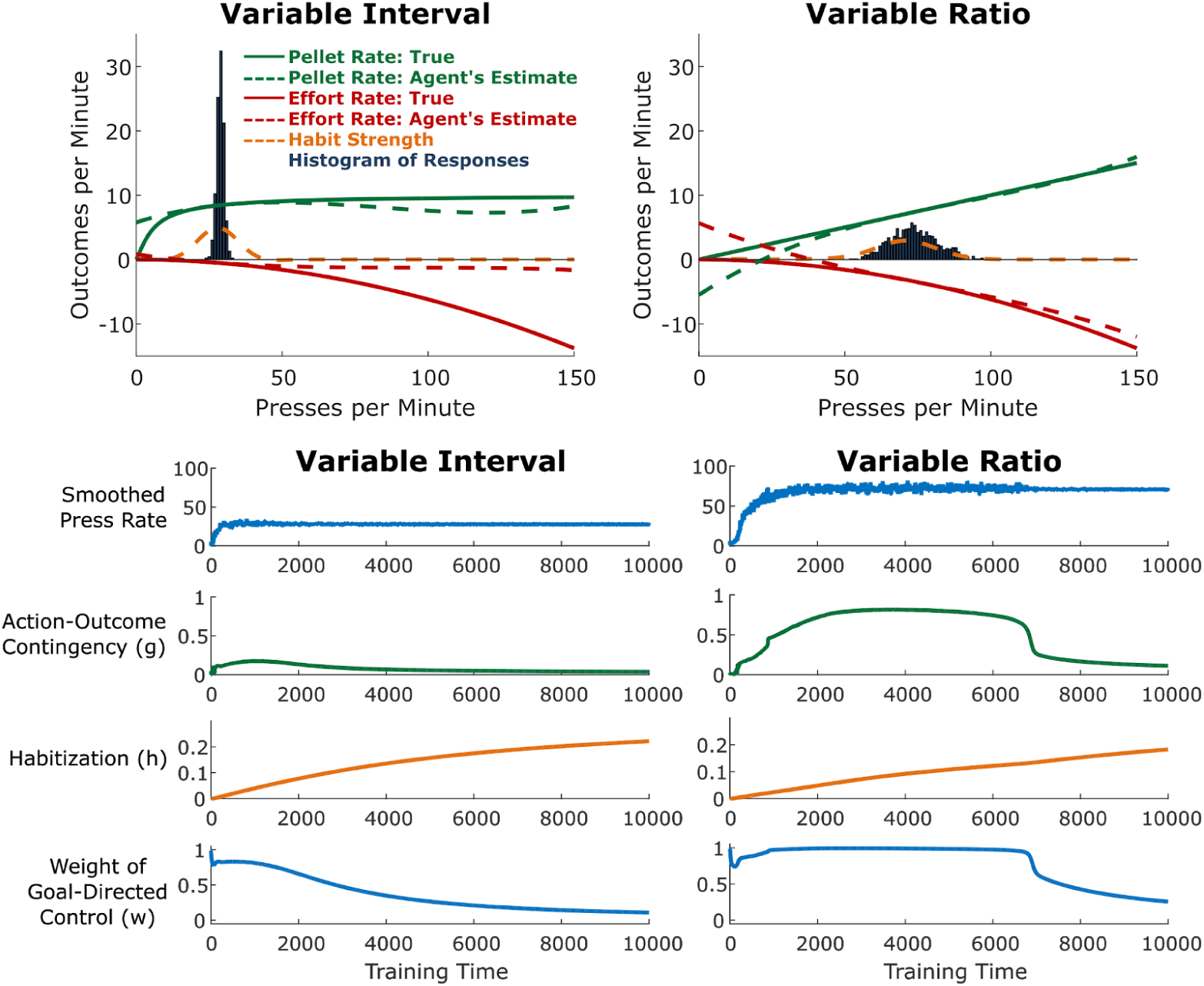
Variable-Interval (VI) schedules produce more rapid habit formation than Variable-Ratio (VR) schedules. Top: Cross-sections of the state of the agent acquiring lever pressing on a VI (left) or VR (right) schedule, taken 5,000 trials into training. Solid curves indicate the rate of pellets or effort as a function of the rate of pressing. Note that in the VR schedule, pellet rate is linear in press rate, whereas in the VI schedule, the relationship is sublinear. Dashed red and green curves indicate the goal-directed system’s estimates of these quantities (*R*). The dashed orange curve indicates the habit strength (*H*) associated with each press rate. Bars give a histogram of the responses of the agent between time points 4,000 and 5,000. Bottom: Time courses of key model variables over the course of training.

**Figure 6.**
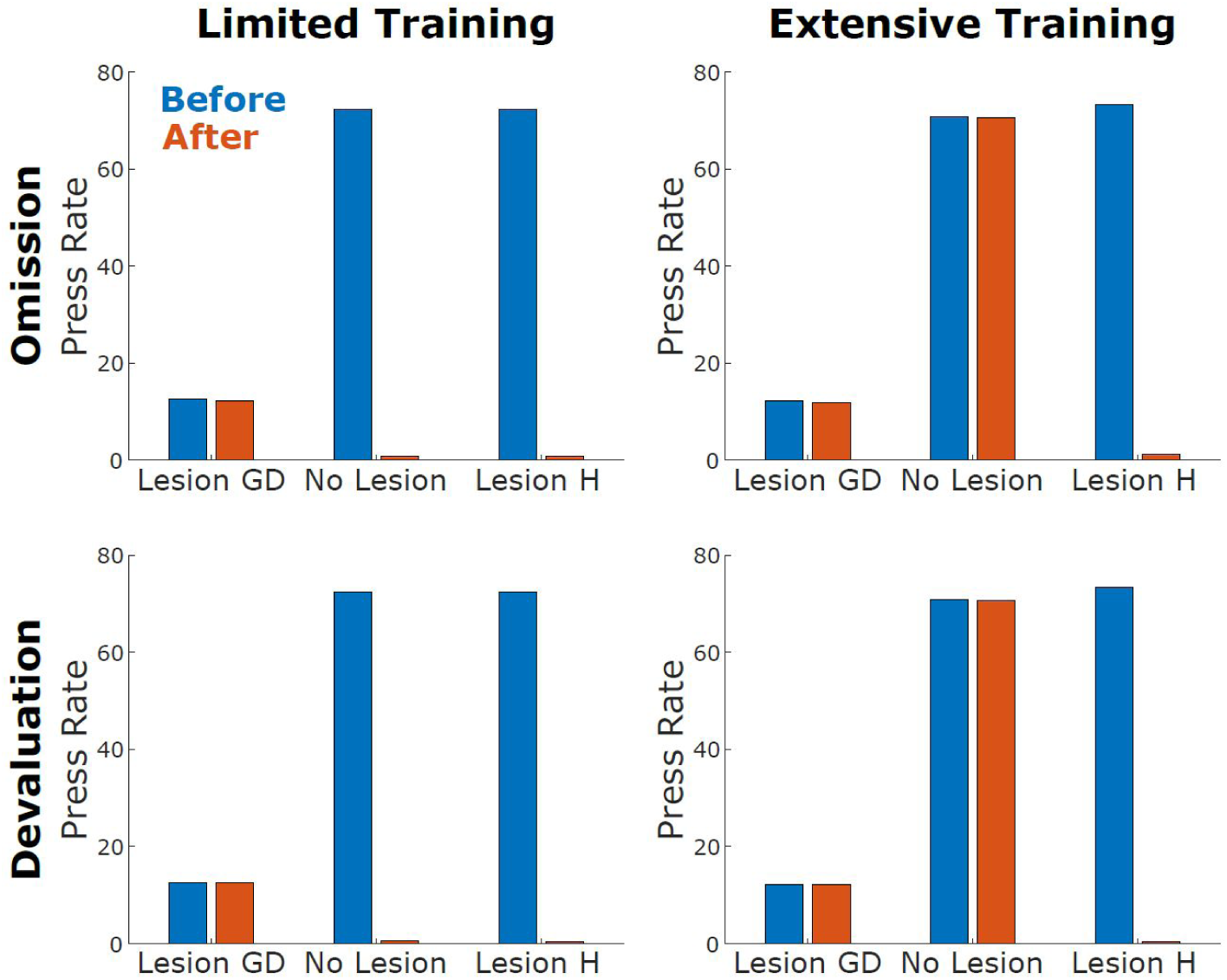
Model Reproduces Effects of Lesions on Behavioral Flexibility. Rate of lever pressing before (blue) and after (orange) omission (top) or devaluation manipulations (bottom rows) performed following either limited or extensive training (left and right columns). We simulated lesions by impairing the goal-directed or habitual controllers, respectively (see Methods for details). The unlesioned model responded flexibly to both manipulations following limited, but not extensive training. Goal-directed lesions caused the model to acquire lever pressing at a much lower rate, and rendered it inflexible to all manipulations, a pattern seen in rats with DMS lesions (Yin et al., 2005). Habit lesions caused the model to respond flexibly to all manipulations, a patterns seen in rats with DLS lesions (Yin et al., 2004, 2006).

### Perseverative Behavior in Sequential Choice Tasks

Finally, we turn to the ubiquitous and poorly understood phenomenon of perseveration, which we argue can be understood as a manifestation of habitual control. In tasks where humans and animals make repeated decisions between similar alternatives, a near-universal observation is a tendency to select actions that have frequently been selected in the past, regardless of their outcome or of the task stimuli. For instance, in instructed task settings with human subjects, the speed and accuracy of an action are enhanced when that action has been recently performed (Bertelson, 1965; Cho et al., 2002). Similar effects are seen in monkeys (Gore et al., 2002). They are also seen in difficult perceptual decision tasks, in which decisions are nominally driven by stimuli that vary from trial to trial in a random way – these effects span monkeys (Gold et al., 2008), rats (Scott, Constantinople, Erlich, Tank, & Brody, 2015), and humans (Akaishi et al., 2014). Perseveration in reward-guided tasks has been seen with the aid of trial-by-trial analyses in rats (Ito & Doya, 2009; Kim et al., 2009), monkeys (Balcarras et al., 2016; Lau & Glimcher, 2005; D. Lee et al., 2005), and humans (Rutledge, Dean, Caplin, & Glimcher, 2010).

In one recent example, rats performing a dynamic two-armed bandit task exhibited behavioral patterns consistent with both reinforcement-seeking (i.e., being more likely to select a recently reinforced action), as well as with choice perseveration (i.e., being more likely to select a recently chosen action). Figure 7 shows the time course of this sensitivity to recent reinforcers (left panel) and recent choices (middle panel) in one example rat (data from Miller et al., in prep). We simulated performance in such an environment and found that the model was able to simultaneously reproduce both the reinforcement-seeking (value-based) and perseverative (value-free) components of these behaviors (Figure 7, right, blue points). Replacing the habitual component of our model with a model-free reinforcement learning system rendered it unable to reproduce the perseverative pattern (Figure 7, right, red points). This example not only begins to validate the predictive abilities of our particular model, but also highlights the importance of a value-free habitual controller more generally in explaining habit-like behaviors that cannot otherwise be accounted for by a model-free RL-based algorithm alone.

**Figure 7.**
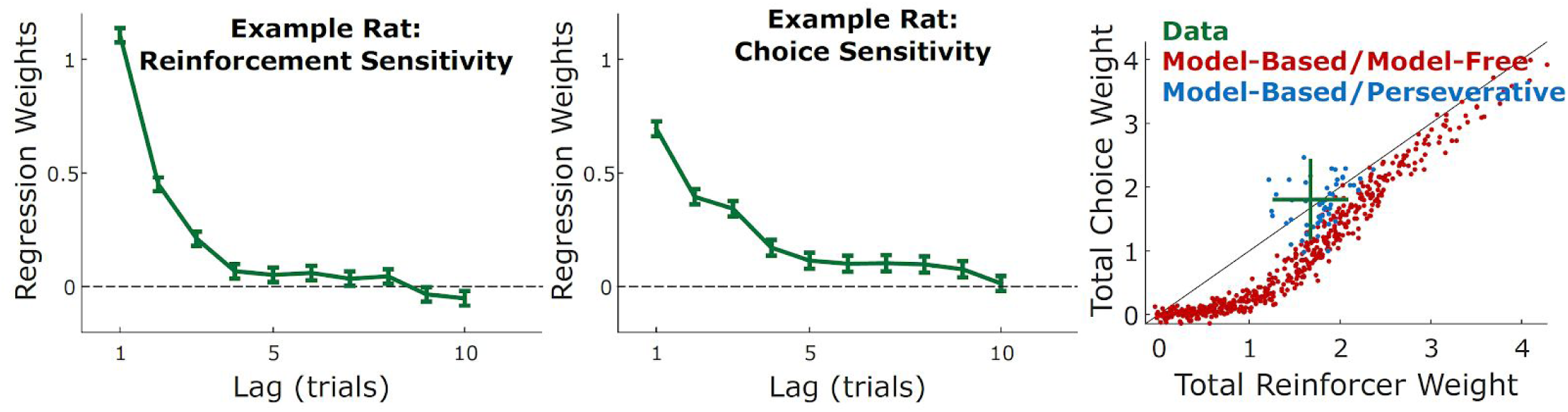
**Left/middle:** Rats performing a sequential choice task exhibit both reinforcer-seeking behavior (left) as well as repetition of recently chosen actions (middle), as has been observed in other species. Reinforcement and choice sensitivity are shown as a function of trial lag for one example rat (Example taken from Miller et al., in prep.). **Right:** To compare the ability of our model and a MB/MF agent to capture key tendencies in these data, we show total reinforcement and choice sensitivity (summing over trial lags shown in left/middle panels) for these rats (green; mean and standard deviation) as well as for simulated model-based/perseverative agents and model-based/model-free agents. Overall the rats exhibit similar choice and reinforcement sensitivity on average. Our model is able to capture this with a relatively limited parameter range (blue scatter; see Table Two); across a much broader parameter range, however, we find that MB/MF agents are unable to generate this same pattern of behavior (red scatter).

## Discussion

Habits are classically thought of as simple, value-free, associations between a situation and the actions most commonly performed in that situation (Dickinson, 1985; Hull, 1943; James, 1890), an intuition that continues to pervade a great deal of modern theorizing (Wood & Neal, 2007; Wood & Rünger, 2016). Despite this legacy, popular computational models of habits hold that they are implemented by value-based mechanisms, learning the expected future reward associated with each action in each situation (Daw et al., 2005; Dolan & Dayan, 2013; Keramati, Dezfouli, & Piray, 2011; S. W. Lee, Shimojo, & O’Doherty, 2014). Here, we have shown computationally that such value-based mechanisms are not strictly necessary, and that a value-free mechanism can account for the major behavioral phenomena that define habits.

We have constructed a computational model in which habits consist of value-free associations between stimuli and actions, and in which these associations are strengthened each time that action is performed in response to that stimulus. This model reproduces key features of the behavioral literature on habits. The first of these features is that habits form slowly over time and often depend on behaviors that are initially taken under goal-directed control. In situations where goal-directed control consistently produces the same behavior in response to the same stimulus, that behavior is likely to become a habit. Once a habit has formed, behavior can become inflexible in the face of changes to the environment that render it no longer desirable, such as contingency omission (in which the reinforcer that drove initial acquisition of the behavior is delivered only when the behavior is *not* performed) and outcome devaluation (in which the reinforcer is rendered no longer valuable to the subject). When combined with another classic idea from the literature on habits – that large action-outcome contingencies delay habit formation – our model is able to explain another classic finding in the literature on habitual control: the effect of reinforcement schedule on the rate of habit formation. Additionally, the proposal that value-free stimulus-response associations exist in the brain explains the ubiquitous observation that human and animal subjects show reinforcer-independent perseverative behaviors in a wide variety of tasks.

This computational account in which habits are understood as value-free stimulus-response associations therefore provides a closer match to classic psychological theories of habits, an account for classic behavioral data on habit formation, and a novel framework for understanding additional behavioral phenomena. As we will describe in the next section, such a mechanism is also more consistent with findings on the neural basis for habitual behavior, and would in turn help to resolve tensions that have emerged in interpreting those findings through the lens of model-free RL.

### Tensions in Neuroscientific Data

#### Separable Neural Substrates for Habits vs. Goal-directed Control

The idea that separate goal-directed and habitual controllers exist in the brain, supported by distinct neural circuits, is strongly supported by lesion data from both humans and other animals. In particular, goal-directed behavior can be disturbed by perturbations to any of a network of interconnected brain regions, including prelimbic cortex (PL; Balleine & Dickinson, 1998; Corbit & Balleine, 2003; Killcross & Coutureau, 2003), dorsomedial striatum (DMS; Yin et al., 2005), mediodorsal thalamus (Corbit, Muir, & Balleine, 2003), basolateral amygdala (Balleine, Killcross, & Dickinson, 2003), and orbitofrontal cortex (Jones et al., 2012; OFC; McDannald, Lucantonio, Burke, Niv, & Schoenbaum, 2011; K. J. Miller, Botvinick, & Brody, 2017). Habitual behavior, on the other hand, can be disturbed by perturbations to infralimbic cortex (Coutureau & Killcross, 2003) as well as the dorsolateral striatum (DLS; Yin et al., 2004, 2006). In human subjects, comparable data are more sparse, but impaired goal-directed behavior has been found in subjects with lesions to ventromedial prefrontal cortex (vmPFC), a candidate homolog of rodent OFC (Reber et al., 2017), as well as following perturbations to dorsolateral prefrontal cortex (dlPFC), a possible homolog of PL (Smittenaar, FitzGerald, Romei, Wright, & Dolan, 2013).

Further support for this idea comes from data measuring neural activity. In rodents, goal-directed behavior results in greater activity in OFC and DMS, while habitual behavior results in greater activity in DLS (Gremel & Costa, 2013). In human subjects, goal-directed value signals during outcome devaluation have been identified in vmPFC (Valentin, Dickinson, & O’Doherty, 2007), and similar signals during contingency degradation have been identified both in vmPFC and in the caudate nucleus, a homolog of DMS (Tanaka et al., 2008). Activity in the putamen, a homologue of rodent DLS, has been found to track the behavioral development of habits (Tricomi, Balleine, & O’Doherty, 2009).

In sum, considerable evidence supports the idea that anatomically separate goal-directed and habitual controllers exist in the brain, and that either controller can be responsible for a given action. Work in rodents points to a number of structures that are necessary for the operation of each of these systems, while work using human subjects, though limited, suggests that the prefrontal and striatal components (at least) are preserved across species (Balleine & O’Doherty, 2010; Liljeholm & O’Doherty, 2012).

#### No Clear Separation for Model-Free vs. Model-Based Control

In contrast to the literature on the habitual/goal-directed dichotomy, such clean dissociations have largely evaded investigations into the neural substrates of model-based and model-free computations, which can theoretically be differentiated in several ways (Doll et al., 2012). The first of these is based on a neuron’s response to an action’s outcome: whereas activity in model-free circuits should only reflect actual reinforcement received (or omitted) and/or the degree to which this deviates from similarly constrained expectations (e.g., temporal difference-based prediction error), activity in model-based circuits should (also) reflect hypothetical (cf. counterfactual/fictive) outcomes that could have been obtained, and should reflect prediction errors based on a richer set of expectations that incorporates, for instance, information about state transition probabilities.

In both of these cases, researchers have been unable to identify circuits that carry uniquely model-based value signals (Bornstein & Daw, 2011; Doll et al., 2012; D. Lee, Seo, & Jung, 2012; Shohamy, 2011). Rather, regions that respond to hypothetical outcomes (a model-based construct) – such as the OFC, vmPFC, and dorsal ACC – tend also to respond to actual outcomes (Abe, Seo, & Lee, 2011; Camille et al., 2004; Coricelli et al., 2005; Hayden, Pearson, & Platt, 2011; Lohrenz, McCabe, Camerer, & Montague, 2007; Rushworth, Noonan, Boorman, Walton, & Behrens, 2011; Strait, Blanchard, & Hayden, 2014). Regions that respond to model-free prediction errors and/or model-free representations of expected value – such as ventral striatum, vmPFC, and even dopaminergic midbrain – also respond to their model-based analogs (Bromberg-Martin, Matsumoto, Hong, & Hikosaka, 2010; Daw, Gershman, Seymour, Dayan, & Dolan, 2011; Kishida et al., 2016; Wimmer, Daw, & Shohamy, 2012). Moreover, ventral striatum also displays signatures of value “preplay” or the covert expectation of reward (Redish, 2016; van der Meer & Redish, 2009), reflective of a classically model-based computation that has been observed in the hippocampus as well. While there are a few notable exceptions to the neuroimaging patterns above – studies that implicate separate regions of striatum in model-free versus model-based valuation (S. W. Lee et al., 2014; Wunderlich, Dayan, & Dolan, 2012) –these studies have not explicitly teased apart model-free valuation from forms of perseveration, leaving open the possibility that model-free value signals in those studies served as proxies for value-free signals of habit strength.

Historically, some of the strongest support for the idea of uniquely model-free computations in the brain has come from studies showing that activity in midbrain dopamine neurons exhibits key characteristics of a computational signal which plays a key role in many model-free learning algorithms: the temporal-difference reward prediction error (Schultz, Dayan, & Montague, 1997). More recent data, however, suggest that dopamine neurons likely carry model-based information as well. In a reversal learning task, these neurons carry information consistent with model-based inference (Bromberg-Martin et al., 2010), while dopamine release in human subjects encodes information about both real and counterfactual reinforcement (Kishida et al., 2016). Perhaps most tellingly, dopamine neurons in a sensory preconditioning task encode prediction errors indicative of knowledge only a model-based system is expected to have (Sadacca, Jones, & Schoenbaum, 2016). Patients with Parkinson’s Disease, in which dopamine neurons die in large numbers, show both impaired model-based behavior (Sharp, Foerde, Daw, & Shohamy, 2015) and increased perseveration (Rutledge et al., 2009), both of which are mitigated by dopamine-restoring drugs. These data indicate that dopamine neurons are unlikely to play a role in a uniquely model-free control system, but instead have access to model-based information and play a role in model-based control.

Collectively, these findings are at odds with the idea that the brain contains separable model-based and model-free systems. Instead, they suggest that to the extent that model-free computations exist in the brain, they are intimately integrated with model-based control, consistent with some existing computational models (Gershman, Markman, & Otto, 2014; Ludvig, Mirian, Kehoe, & Sutton, 2017; Pezzulo, van der Meer, Lansink, & Pennartz, 2014; Silver, Sutton, & Müller, 2008; Sutton, 1990). This lack of clear dissociation between model-based and model-free computations stands in stark contrast to the dissociations (described above) between circuits for goal-directed and for habitual control. This casts doubt on the idea that a one-to-one mapping exists between these two dichotomies, and motivates the search for alternative accounts (K. J. Miller, Shenhav, Pezzulo, & Ludvig, 2018).

### A Proposed Realignment

To overcome the obstacles just described, we propose a revised framework with two key alterations: a divorce and a union. We first propose severing the tie between habits and model-free RL, and instead defining a category of behaviors that are “value-free” and therefore distinct from either type of RL computation. These behaviors would consist of S-R associations whose strengths are modified primarily through repetition, consistent with Thorndike’s Law of Exercise and some more contemporary notions of habit learning (e.g. direct cuing; Wood & Rünger, 2016). They would be value-free in the sense that such a behavior could be fully described from the triggering of the stimulus to the emission of a response without requiring the representation of expected reinforcement along the way. Being value-free, however, would not entirely prevent these behaviors from being sensitive to one’s surroundings. S-R associations can be learned in a fashion such that their likelihood of being triggered is influenced by the spatial, temporal, and motivational contexts. Moreover, and perhaps counterintuitively, being value-free would by no means prevent S-R associations from being *value-sensitive*. In particular, while the mechanism for S-R learning might be through repetition, the strength of the resulting association might be influenced by the value of the action being repeated. That is, the behavior that becomes habitual may initially have been performed, and gradually strengthened, while in the pursuit of value under the goal-directed controller, but once a habit has formed, behavior is no longer directly driven by value (Tricomi et al., 2009; Wood & Rünger, 2016).

Our second proposal is to re-unify model-free and model-based computations as being two different drivers of goal-directed (i.e., value-based) behavior, distinct from the class of value-free behaviors just described. Rather than viewing these two computations as categorically distinct, we further suggest that it may be more appropriate to view them as falling along a continuum, varying according to the amount of information used to make decisions. On this view, the available information would range from recent rewards, through simple relationships between stimuli, up to a full world model of all possible states. All of these computations are goal-directed, but their informational content directs them toward different goals. This latter proposal carries an additional benefit in that it obviates the need to cache value in a “common currency” (i.e., without reference to a specific outcome like juice or food type). Storage of such a common currency signal is typically required for model-free RL, but evidence for such signals in the brain remains weak (Morrison & Nicola, 2014; O’Doherty, 2014; Schoenbaum, Takahashi, Liu, & McDannald, 2011). This realignment therefore offers the possibility of bypassing model-free RL computations entirely, but our current model is agnostic as to whether such a drastic revision is appropriate based on the available evidence.

### Relationship to Previous Computational Models

A large and influential body of computational work is built on the assumption that habitual control arises from model-free reinforcement learning algorithms (Dolan & Dayan, 2013; O’Doherty, Lee, & McNamee, 2015). This work originates from a proposal by Daw and colleagues (Daw et al., 2005) that the parallel goal-directed and habitual controllers described by animal learning theory can be understood computationally as model-based and model-free reinforcement learning agents, operating in parallel and competing with one another for control of behavior. Subsequent work in this line has proposed different mechanisms for this competition (Keramati et al., 2011), or suggested ways in which model-based and model-free controllers might cooperate rather than compete (Keramati, Smittenaar, Dolan, & Dayan, 2016; S. W. Lee et al., 2014), but has retained the basic premise that habits are instantiated by model-free RL mechanisms and that habitization can be understood as a process by which these mechanisms come to dominate model-based mechanisms for the control of behavior.

This view of habitization is most successful at explaining the inflexibility of habits in the face of outcome devaluation: because model-free mechanisms associate actions with common-currency values only, rather than particular outcomes, they are unable to respond flexibly when a particular outcome is no longer desired. This view is in tension, however, with the inflexibility of habits in the face of contingency omission: learning that an action which previously led to reinforcement now instead prevents reinforcement should be well within the capabilities of a model-free system. The only resolution to this tension that we are aware of requires invoking an additional habitization mechanism: a slow decrease in the learning rate of the model-free system when faced with stable environments (Dayan, Kakade, & Montague, 2000). Our proposal avoids this tension entirely by positing that habits are instantiated not by model-free RL, but by mechanisms that are entirely value-free. It therefore explains the inflexibility of habitual behavior in the face of both devaluation and omission using only one mechanism: the handoff of control from a value-based to a value-free system.

In a similar vein to the current proposal, Dezfouli and Balleine (2012) developed a model of habits that dropped the mapping between habits and model-free reinforcement learning. Instead, they proposed that habits should be modeled as learned action sequences (“chunks”). In contrast to our model, however, those action sequences are initiated under (outcome-sensitive) goal-directed control, after which they proceed in an outcome-insensitive manner until the sequence is completed. A particular sequence of actions can be executed more quickly when selected as a chunk than when each action is selected individually in series, and this strategy is preferred when the benefit of speeded responses outweighs the cost of such temporarily open-loop control. In contrast, the model of habits we are proposing completely cuts the tie between reinforcement learning and the habitual controller. Actions become habitized merely from use in a particular state, independent of any costs or benefits. This view provides an alternative explanation for the observation of habitized action sequences: when actions are typically performed in a particular order, the proprioceptive or other feedback associated with each action can become the “stimulus” that directly cues the subsequent action in the sequence (see James, 1890, chapter 4). Exploring this idea using using computational RL algorithms would involve building environments in which information about the previous action is incorporated into the state space. Constructing such models, and designing experiments to dissociate them from the Dezfouli and Balleine account, is a promising direction for research into habit formation. Such experiments might involve interposing additional instructed actions into traditional sequential behavior assays of habit formation.

A mechanism for the development of automaticity that is conceptually similar to ours has appeared in models of interactions between the cortex and the basal ganglia (Ashby, Turner, & Horvitz, 2010). These models propose that novel behaviors are first acquired via a dopamine-dependent plasticity mechanism within the basal ganglia, and that with consistent performance, control of behavior is transferred to cortex via a Hebbian cortico-cortical plasticity mechanism. Developed first in the context of categorization learning (Ashby, Ennis, & Spiering, 2007), this idea has recently been applied to sequence learning (Hélie, Roeder, Vucovich, Rünger, & Ashby, 2015) and to action selection in probabilistic environments (Topalidou, Kase, Boraud, & Rougier, 2017), and it has been suggested to describe how basal-ganglia-dependent behavior becomes automatic in general (Hélie, Ell, & Ashby, 2015). Though the cortical module of these models is conceptually similar to our habitual controller, the basal ganglia module is different from our goal-directed controller in important ways: its learning rule instantiates a version of model-free RL, which tends to repeat actions in situations where they have led to reinforcement in the past, but does not learn about the particular outcomes that are expected to follow each action. Such a mechanism is not expected to exhibit the critical properties that characterize goal-directed control, most notably flexibility in the face of outcome devaluation. This form of flexible behavior is thought to require model-based mechanisms (Daw et al., 2005).

In addition to utilizing different mechanisms for value-based control and for arbitrating between the value-based and value-free controllers, our model also differs from these corticostriatal models in the level of analysis at which it is described. While this limits the level of detail with which our model can engage neurobiological data, it greatly facilitates engagement with a wide variety of behaviors and with a broad range of other theoretical approaches (Frank, 2015; Frank & Badre, 2015). As such, we have applied the model to a wider cut of behaviours, including outcome devaluation, perseveration, and the impact of reinforcement schedule on habitization. A critical next step, however, will be to develop a neurobiologically detailed implementation of our competing controllers, building on the types of multiple learning systems described above and related work (Ashby, Alfonso-Reese, Turken, & Waldron, 1998; McClelland, McNaughton, & O’Reilly, 1995; O’Reilly & Frank, 2006). Such work would seek to integrate the Hebbian learning systems of these earlier models with a neurobiologically plausible model-based controller (Friedrich & Lengyel, 2016; Solway & Botvinick, 2012)

Many formalizations of the standard mapping from habits/goals to model-free/model-based reinforcement learning also include a perseveration kernel (e.g. Daw et al., 2011; Lau & Glimcher, 2005). That is, in addition to the two types of value learning, subsequent choice is also influenced by the most recent choice. A similar tendency to repeat actions also appears due to predictive coding in the free energy framework, whereby actions are repeated for maximal predictability (Pezzulo, Rigoli, & Friston, 2015). This extra piece of computational machinery allows the models to account for the tendency to repeat choices, independent of values. Here, we bring this perseveration kernel to the foreground. Our proposed framework re-maps habits to the perseveration kernel and provides an account of how that kernel might plausibly operate in tandem with a goal-directed controller so as to account for behaviors that have previously been described by RL models. In effect, we are showing that the model-free component of some of these previous formalizations might not be necessary and that an elaboration of this perseveration kernel actually serves as a better model of habits.

In our model, control of behavior is allocated on each trial to either the habitual or the goal-directed system by an arbiter. This arbiter is similar to mechanisms found in computational accounts mapping habitual/goal-directed control onto model-free/model-based reinforcement learning. In the initial formalization of this idea (Daw et al., 2005), each system determined the uncertainty in its value estimates, and the arbiter selected the system with less overall uncertainty. The extra computational complexity associated with the goal-directed system was a source of uncertainty, leading the arbiter to favor the habitual system in well-learned environments. Subsequent accounts have developed arbiters that consider proxies for this uncertainty (S. W. Lee et al., 2014), or other possible advantages of habitual over goal-directed control, such as response time (Keramati et al., 2011). The arbiter in our model is motivated by a classic observation in research on habits: habits are promoted in situations where the contingencies between actions and outcomes are weak (Dickinson, 1985). Future research should explore the conditions under which these arbiters make similar or diverging predictions for habitual control and systematically test the relative success of these arbitration approaches at accounting for empirical data under those conditions.

### Implications

The realignment we are proposing carries important implications and testable predictions for future work. First and foremost, our account predicts that neural circuits associated with habitual behavior (e.g., DLS) should also be related to (value-free) perseveration. We might therefore expect greater activity in this circuit with additional repetitions of a previous action, and that lesioning parts of this circuit will reduce the tendency to perseverate. Secondly, we predict that elicitation of action repetition should be sufficient to construct new habits, without requiring reinforcement. For instance, generating actions with microstimulation in a particular context may facilitate the subsequent performance of those actions in that same context. Such evidence would provide strong support for our model. This prediction also provides a mechanistic underpinning for the repetition strategies that have shown to be effective at improving workplace and health-related performance through habit formation (Gardner, Lally, & Wardle, 2012; Lally et al., 2010; Wood & Rünger, 2016). Related to both of these claims, our model suggests that disorders of habitual behavior (e.g., Obsessive-Compulsive Disorder, Tic Disorders) need not result from dysfunction in valuation (cf. Gillan & Robbins, 2014). Our model can help to tease apart the degree to which value-free versus value-based processes are implicated in each of these disorders, and this will have important implications for considerations of etiology and treatment. Our model makes additional but weaker predictions with respect to the relationship between model-free and model-based processes. If these represent related manifestations of a common value-based system, we expect brain regions that reflect model-free value signals to also reflect model-based value signals, as has been the case in many previous studies (Abe et al., 2011; Bornstein & Daw, 2011; Doll et al., 2012; D. Lee et al., 2012; Shohamy, 2011). For instance, model-free prediction errors should not be found in regions that fail to exhibit model-based prediction errors. Related to this, one should not be able to lesion part of the model-free valuation circuit without influencing model-based behavior. To the extent that model-based forms of decision-making draw on additional mechanisms, including hippocampally-mediated stimulus-stimulus associations (Bornstein & Daw, 2013; Bunsey & Eichenbaum, 1996; Dusek & Eichenbaum, 1997) and prefrontal control mechanisms (E. K. Miller & Cohen, 2001), the reverse need not be true; we would predict that inactivating (K. J. Miller et al., 2017; Smittenaar et al., 2013) or otherwise drawing resources away from (Otto, Gershman, Markman, & Daw, 2013; Otto, Skatova, Madlon-Kay, & Daw, 2015) such additional mechanisms would selectively impair model-based behavior, as has been observed. Thus, our model accommodates data that fail to identify a model-free learning component in behavior and/or neural activity, while also accommodating a growing literature demonstrating factors that selectively promote or inhibit model-based control.

Importantly, the value-free mechanisms we have proposed for habits by no means preclude a role for value or motivational state (e.g., hunger or satiety) in habit learning. These may modulate the strengthening of habit associations either directly (e.g., through a feedback mechanism that influences the S-R association) or indirectly (e.g., by influencing the vigor of an action, which in turn results in greater associative strengths). The particular form of such a mechanism that best accounts for available data is a matter of further research, and one which we aim to pursue in extending our model.

## Conclusions

We have provided evidence that a value-free learning process -- according to which S-R associations are strengthened through action repetition in a Hebbian manner -- may be sufficient to generate behaviors that have been traditionally classified as habits, and held up in contrast to goal-directed behaviors. We demonstrate that such a mechanism leads to perseveration of a previously higher-value action following contingency degradation or outcome devaluation and increased perseveration of all actions in a probabilistic choice task with varying action-outcome contingencies. We further show that such habitual behaviors are diminished by simulating lesions to a habitual system, consistent with classic findings in the animal behavior literature. Crucially, the system that generates these habitual behaviors does so without engaging in any manner of reinforcement learning (model-free or otherwise), consistent with theories that place habits outside the domain of RL.

In spite of the absence of a model-free controller, we also show that our model can still capture key features of behavior for a task that is thought to rely on both model-free and model-based control, bringing into sharper focus recurring questions about whether and/or when a purely model-free controller is necessary to explain these behaviors. Collectively, we argue that these findings support a realignment of current computational models of decision-making, towards (re-)associating goal-directed/habitual with value-based/value-free rather than model-based/model-free. Beyond providing a potentially more parsimonious account of previous behavioral results, such a realignment may offer a better account of extant neural findings, including the fact that structures associated with model-free and model-based computations (i.e., value-based computations) tend to overlap, whereas lesion/inactivation studies have revealed clear dissociations between structures associated with goal-directed behavior versus (potentially value-free) habits.

## Acknowledgements

We would like to thank Nathaniel Daw, Yael Niv, Matthew Botvinick, Jonathan Cohen, Charles Kopec, Kimberly Stachenfeld, Oliver Vikbladh, Marcelo Mattar, and Sebastiaan van Opheusden for helpful discussions.

## Appendix A: Model for Environments with Multiple States

This appendix provides the complete equations for the full version of the model that can include multiple states. For simplicity, the version in the text only includes a single state because the simulations here all include only a single state.

### Habitual Controller

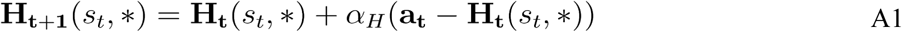

where *s*_*t*_ is the current state, **H**_**t**_(s_t_,*) is the row of **H**_**t**_ corresponding to *s*_*t*_, *α*_*H*_ is a step-size parameter that determines the rate of change, and **a**_***t***_ is a row vector over actions in which all elements are zero except for the one corresponding to *a*_*t*_, the action taken on trial *t*. Importantly only the row of **H** corresponding to *s*_*t*_ is updated – other rows remain the same.

### Goal-Directed Controller

The goal-directed controller maintains an estimate, **R**_**t**_ of predicted immediate reinforcement, in which *R*_*t*_*(s,a,m)* gives the agent’s expectation at timepoint *t* of the magnitude of reinforcer type *m*, that will follow from action *a*, in state *s*. Initial reinforcement expectation **R**_**0**_ is set to zero, and after each trial, the agent updates these quantities according to the following equation (Sutton & Barto, 1998):

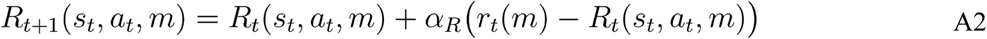

where *s*_*t*_ is the current state, *a*_*t*_ is the current action, *r*_*t*_*(m)* is the magnitude of the reinforcer of type *m* received following that action, and *α*_*R*_ is a step-size parameter which governs the rate of learning.

The goal-directed agent also maintains an estimate **T** of transition probabilities, where each element, *T*_*t*_*(s,a,s’)*, gives the agent’s expectation at timepoint *t* that taking action *a* in state *s* will lead to subsequent state *s’*To ensure proper normalization, **T**_**0**_ is initialized to *1/n*, where *n* is the total number of states in the environment and is updated according to:

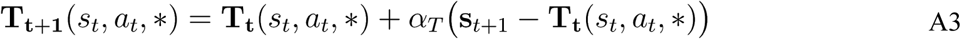

where **T**_**t**_(s_t_,a_t_,*) is the slice of **T**_**t**_ corresponding to *s*_*t*_ and *a*_*t*,_, and **s**_**t+1**_ is a row vector over states in which all elements are zero except for the one corresponding to *s*_*t+1*_, the state visited on trial *t*, and *α*_*T*_ is a step-size parameter. The goal-directed agent assigns values to states based both on expected immediate reinforcement as well as on expected reinforcement in future states via the recursive Bellman equation (Sutton & Barto, 1998):

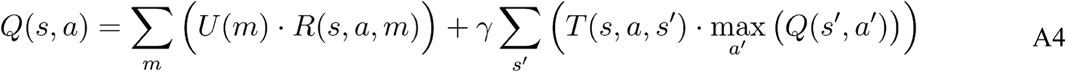

where *U(m)* is a utility function giving the value that the agent assigns to reinforcers of each type *m*, and γ is a rate of temporal discounting giving the relative utility of immediate vs. future rewards.

### Arbiter

The arbiter governs the relative influence of each controller on each trial. It computes an overall drive *D(s,a)* in favor of each action, *a*, taken in each state *s*, as a weighted sum of the habit strength **H** and the goal-directed value **Q**:

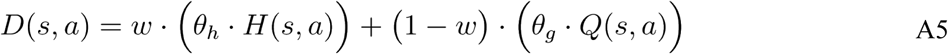

where *θ*_*h*_, and *θ*_*g*_ are scaling parameters, and *w* is a weight computed on each trial by the arbiter to determine the relative influence of each controller (see Equation A9). The model then selects actions according to a softmax on **D**:

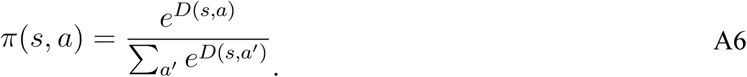

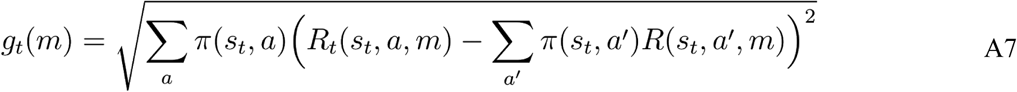

which reflects the degree of variation in expected outcome for that reinforcer, based on the available actions and the policy. The arbiter also computes an analogous quantity for the habitual controller, which we term “overall habitization” *h*:

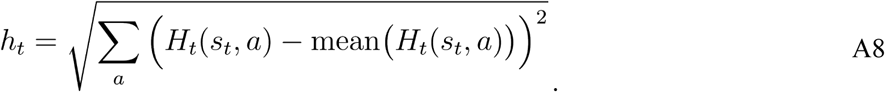

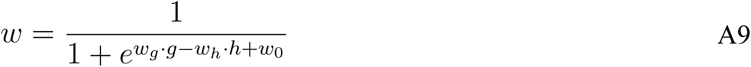

## Appendix B: Model-Based/Model-Free Agents

In simulations of the two-armed bandit environment, we compare our model to one consisting of a mixture of model-based and model-free agents. In a general environment, the model-based agent would be identical to the goal-directed agent described in Appendix A. In the two-armed bandit environment, however, there is only one state and one type of reinforcer, so the equations describing this agent (equations A2, A3, and A4) can be simplified to the following single equation:

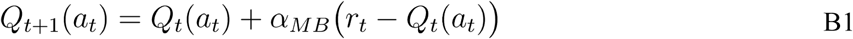

where the parameter α_MB_ governs the rate of learning in this agent. This environment is therefore simple enough that the model-based agent does not employ any of its uniquely model-based machinery. The model-free agent is described by a similar equation, where we use **H** to denote its values, because in model-based/model-free schemes the model-free controller is typically thought of as implementing habitual control (Daw et al., 2005).

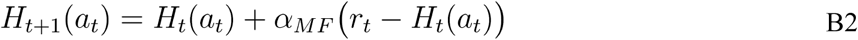

where the parameter α_MF_ governs the rate of learning. The model-based and model-free values are combined to determine overall drive:

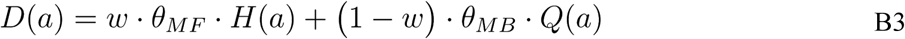

where *θ*_MB_ and *θ*_Mf_ are scaling parameters, and *w* is a mixing weight, and drive is used to determine choice:

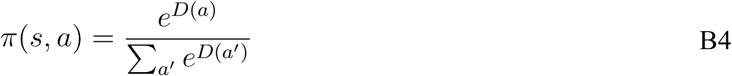

The mixed model-based/model-free controller therefore has five free parameters, α_MB_, α_MF_, *θ*_MB_, *θ*_MF_, and *w*.

